# Transcriptional dynamics of transposable elements in the type I IFN response in *Myotis lucifugus* cells

**DOI:** 10.1101/2022.04.18.488675

**Authors:** Giulia Irene Maria Pasquesi, Conor J. Kelly, Andrea D. Ordonez, Edward B. Chuong

## Abstract

**Background:** Bats are a major reservoir of zoonotic viruses, and there has been growing interest in characterizing bat-specific features of innate immunity and inflammation. Recent studies have revealed bat-specific adaptations affecting interferon (IFN) signaling and IFN-stimulated genes (ISGs), but we still have a limited understanding of the genetic mechanisms that have shaped the evolution of bat immunity. Here we investigated the transcriptional and epigenetic dynamics of transposable elements (TEs) during the type I IFN response in little brown bat (*Myotis lucifugus*) primary embryonic fibroblast cells, using RNA-seq and CUT&RUN.

**Results:** We found multiple bat-specific TEs that undergo both locus-specific and family-level transcriptional induction in response to IFN. Our transcriptome reassembly identified multiple ISGs that have acquired novel exons from bat-specific TEs, including *NLRC5*, *SLNF5* and a previously unannotated isoform of the *IFITM2* gene. We also identified examples of TE-derived regulatory elements, but did not find strong evidence supporting genome-wide epigenetic activation of TEs in response to IFN.

**Conclusion:** Collectively, our study uncovers numerous TE-derived transcripts, proteins, and alternative isoforms that are induced by IFN in *Myotis lucifugus* cells, highlighting candidate loci that may contribute to bat-specific immune function.

## BACKGROUND

Bats are increasingly recognized to be an important reservoir of zoonotic viruses, including Rabies viruses, Dengue viruses, Ebolaviruses, and Coronaviruses [1, 2]. Remarkably, viral infection in bats is associated with minimal lethality and reduced inflammatory phenotypes, which has led to extensive research aimed at uncovering bat-specific features of immunity [3–6].

Recent genomic and functional studies in bats have begun to reveal species-specific adaptations affecting innate immune responses. For example, the interferon (IFN) genes have been subject to evolutionary expansions and contractions in different bat species, and several species exhibit constitutive expression of IFNs at low levels [7] (reviewed in [1]. Bats also exhibit unique subsets of ISGs [8]. A time-course analysis comparing the type I IFN response in *Pteropus alecto* and humans revealed distinct kinetics of IFN-stimulated gene (ISG) regulation, where bats exhibit more rapid downregulation of ISGs compared to humans [9]. In addition to adaptations affecting IFN signaling, other pro-inflammatory genes are also frequently mutated or lost, including TLR genes [10], components of the inflammasome [11, 12], the cGAS/STING pathway [13], and the OAS/RNASEL pathway [9, 14]. These studies have begun to reveal the genetic basis for bat-specific features of immunity which could help us understand their propensity to act as viral reservoirs.

While it is clear that bats have evolved numerous unique adaptations affecting innate immune pathways, we still have a poor understanding of the genetic mechanisms responsible for these changes. Our study focuses on TEs as a potentially important yet understudied source of mutations that shape bat immunity. In studies of other mammalian lineages, there is evidence that lineage-specific TEs contribute to innate immune functions through a variety of mechanisms. For example, TE and virus-derived proteins have been repeatedly co-opted as immune proteins that often restrict viruses through dominant negative activity [15] in ruminants [16], rodents [17], and primates [18]. TE-derived non-coding transcripts can readily form immunostimulatory double-strand RNAs or DNAs [19–21]. Finally, TEs can regulate interferon-inducible gene expression by acting as regulatory elements [22–26]. The recurrent co-option of TEs for immune functions throughout evolution may reflect their capacity to fuel adaptation by increasing genomic variation, especially in the context of host-pathogen coevolutionary arms races [27, 28].

TEs are widely speculated to have been important contributors to the evolution of bats [29–33], including bat-specific immune functions [34, 35]. While the genomes of all mammalian species contain numerous lineage-specific transposons, bat genomes are distinguished by recently active DNA transposons, which are extinct in most other mammalian lineages [29–32]. In addition to DNA transposons, bat genomes have been extensively shaped by other TEs typically found in other mammals, including LTR retrotransposons like endogenous retroviruses, LINEs, and SINEs. Notably, a recent report identified a *Rhinolophus-*specific LTR insertion within an exon of *OAS1,* which disrupts horseshoe bat antiviral activity to SARS-CoV2 with significant implications for the OAS1*-*mediated response to SARS-CoV2 in humans [14]. However, aside from this example, the potential impact of TEs on bat immunity remains largely unstudied, due to the lack of experimental and functional genomic resources available for studying bat immunology.

To conduct a comprehensive genome-wide study of TEs in bat innate immunity, we conducted transcriptomic and epigenomic profiling of the type I IFN response in *Myotis lucifugus* primary fibroblast cells. We used RNA-seq and CUT&RUN to characterize the IFN-inducible transcriptomes and regulatory elements, which allowed us to systematically examine the contribution of TEs to loci that define the bat IFN-inducible response.

## RESULTS

To characterize the contribution of TEs to the IFN response in bats, we conducted transcriptomic and epigenomic profiling of the type I IFN response in *M. lucifugus* primary embryonic fibroblast cells (Fig. 1). We stimulated cells using recombinant universal IFN alpha (IFNa), and profiled the transcriptome at 0, 4 and 24h time points using RNA-seq. We confirmed cellular response to universal IFN treatment using qPCR on canonical ISGs (Fig. S1), as shown previously for *M. lucifugus* dermal fibroblasts [8]. We also profiled 0 and 4h time points using CUT&RUN to map genome-wide localization of H3K27ac, POLR2A, and STAT1. We aligned these data to a chromosome-scale HiC assembly of the little brown bat genome (myoLuc2.0_HiC) [36], which was the most contiguous assembly available (Scaffold N50 of ∼95.5Mb).

**Fig. 1.**
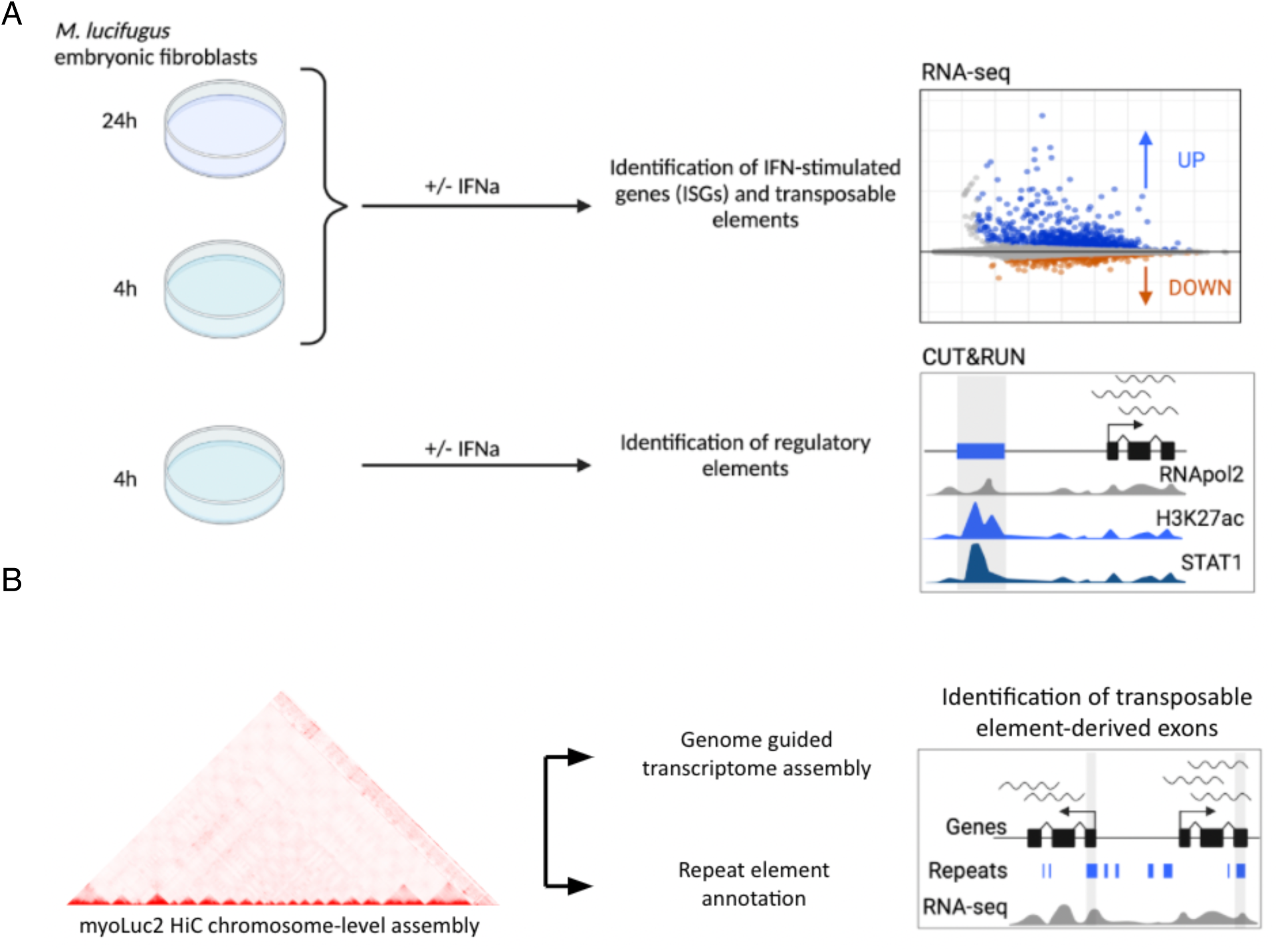
Experimental design. **A**) *Myotis lucifugus* embryonic fibroblast primary cells were treated for 24 hours (24h) and for 4 hours (4h) with 1000U/ml of universal IFNa (+IFNa) or matched volume of DPBS (-IFNa). Total RNA at both time points was extracted and used as input for RNA library preparation and sequencing to identify differentially expressed genes and transposable elements (TEs). To characterize changes in chromatin accessibility upon IFN treatment, cells were similarly treated for 4h, and subjected to the CUT&RUN protocol on H3K27ac, POLR2A, and STAT1. **B**) The chromosome-level genome assembly for *Myotis lucifugus* was used as reference to perform *de novo* repeat element identification and annotation. Combined with genome guided transcriptome assembly of our RNA-seq datasets, the custom repeat element annotation was used to identify TE-derived and virus-derived isoforms and transcription start sites (TSS).

Prior to analyzing our functional genomic data, we performed de-novo repeat identification on the myoLuc2.0_HiC assembly using RepeatModeler2 [37–39]) and HelitronScanner ([37–39], followed by repeat annotation using RepeatMasker [40]. We annotated 42.7% of the genome as derived from TEs, compared to 35.5% annotated in the myoLuc2 non-HiC assembly (https://www.repeatmasker.org/species/myoLuc.html). L1 LINEs represent the most abundant TE group (14.1%), followed by virus-derived elements (ERVs; 5.6%) and DNA hAT and Helitron elements (3.7% and 3.2% of the genome, respectively) (Fig. 2A). As previously identified [29], our analyses show the lineage-specific expansion of DNA transposons, first led by Helitron elements, and more recently by multiple subfamilies of hAT elements that may have been introduced by horizontal transfer (Fig. 2A) [41].

**Fig. 2.**
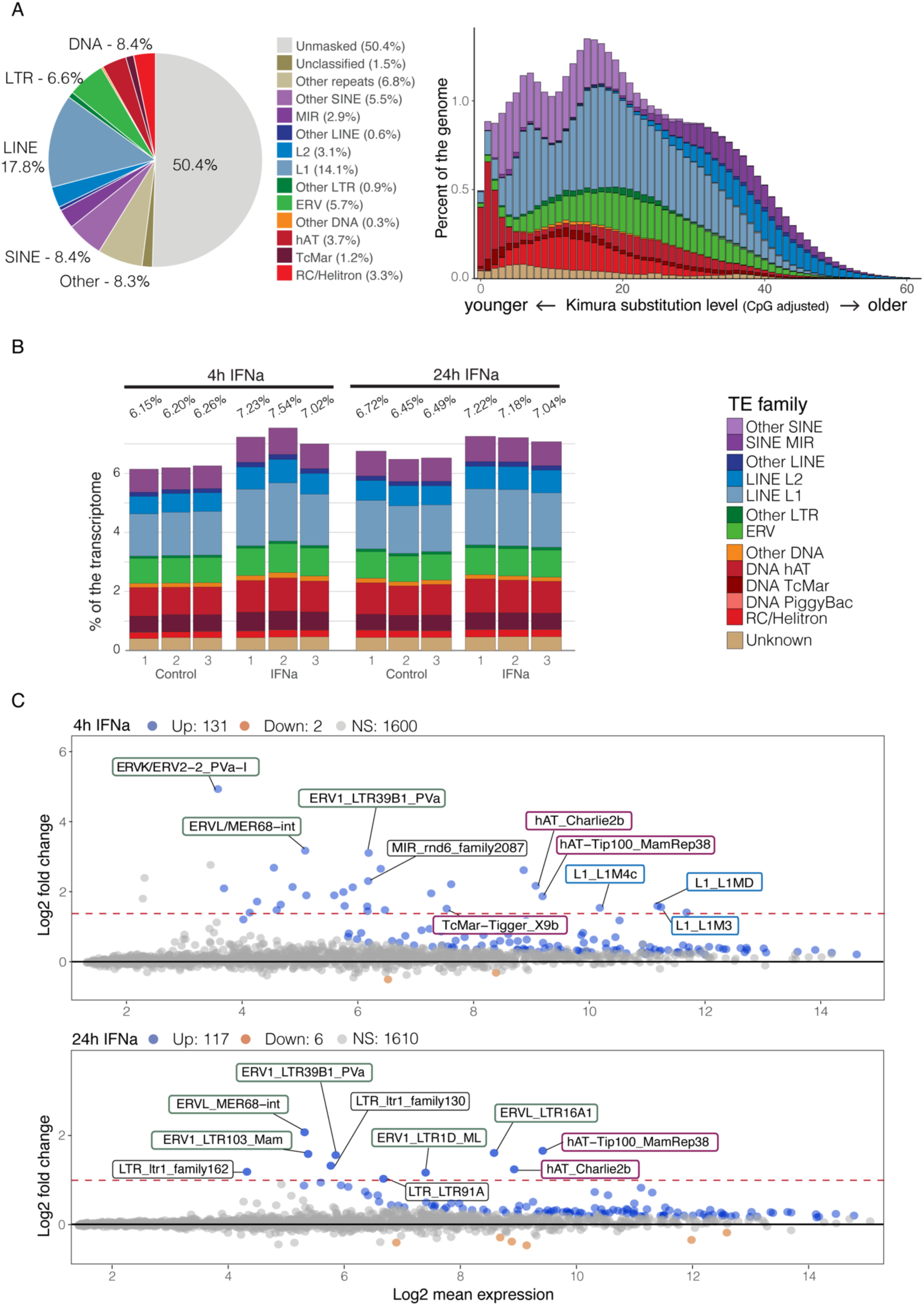
Repeat element composition, evolutionary dynamics and transcriptional profiles. Repeat elements were annotated by combining *de novo* identification and homology-based searches. **A**) Pie chart shows the relative abundance of main TE families (42.7% total) as percentage of the genome; histogram shows the composition as percentage of the genome of major TE superfamilies as a function of the divergence (Kimura2Distance) from the reference consensus sequence of each TE. Given the correlation between divergence from the consensus and time of transposition, the lower the K2D (left) and the younger a TE is. As previously identified in Myotis and other bat species, we recovered recent expansion and activity of multiple DNA elements, in particular Helitrons and more recently hAT elements. **B**) Histograms show expression levels of major TE families as a fraction (%) of normalized read counts from RNA sequencing data of IFNa-stimulated and unstimulated cells at 4h and 24h post treatment. **C**) MAplots of apeglm [83] transformed data show differentially expressed TEs at 4h (top) and 24h (bottom) post IFN treatment. For the 4h time point, only the top three TEs per family with the highest Log2 fold change were labelled. For both time points, only TEs that met a threshold of adjusted p-value < 0.05 and log2 fold change > 1.5 (accepted fdr = 0.05) were labelled. Counts of upregulated (blue dots) and downregulated (orange dots) TEs are based on adjusted p-value (<0.05; accepted fdr = 0.05) only. Among induced TEs at 4h (45 total based on our cutoffs) we recovered for the most part ERV retrotransposons (green outline), DNA hAT transposons (red outline) and L1 LINEs (light blue outline). At 24h post treatment we only found 10 families that met the filtering criteria, for the majority ERV shared with the 4h dataset.

### Gene and TE expression profiles upon IFN stimulation

To analyze transcriptional activity at the TE family level, we mapped RNA-seq reads to both genes and TE families using TETranscripts (Fig. 2B) [42]. On average, 6.38% of RNA-seq reads mapped to TEs in unstimulated cells, while 7.26% of reads mapped to TEs after 4h IFN treatment, and 7.15% after 24h IFN treatment. The most abundant TE-derived transcripts we identified included L1 LINEs, DNA/hAT elements, ERVs, SINEs and L2 LINEs. We identified 45 TEs that showed significant family-level transcriptional induction at 4h (adj. *p*-val < 0.05; log2FC > 1.5), and 8 families induced at 24h according to the same cutoff thresholds (Fig. 2C). These included multiple ERV families (21), L1 LINEs (6) and DNA transposons (6) at 4h post treatment, and ERVs (5) at 24h post IFN treatment. These findings indicate that multiple bat-specific TEs show family-level transcriptional induction in response to IFN treatment, peaking at 4h and diminishing but still present at 24h post induction.

We next analyzed our RNA-seq dataset to identify IFN-stimulated genes (ISGs), as determined by DESeq2 comparing gene counts from treated to untreated samples in pairwise comparisons at 4h and at 24h post treatment (Tables S1 and S2). We first used the homology-based annotation provided by DNAzoo as a reference transcriptome. Using a cutoff of adj. *p*-val < 0.05 and log2FC > 1.5, we identified 213 upregulated transcripts (corresponding to 138 unique genes) and 1 downregulated gene at 4h, and 91 upregulated transcripts (corresponding to 66 unique genes) at 24h post IFN stimulation. Based on their expression dynamics, 4 main transcript clusters were identified (Fig. 3A): I) transcripts showing a strong response to IFN at 4h that declines at 24h; II) transcripts showing mild induction at 4h and decline at 24h; III) transcripts showing mild, stable induction; and IV) transcripts showing strong induction at 4h and rapid decline to levels similar to unstimulated cells at 24h. Both 4h and 24h post-induction ISGs were enriched for canonical ISGs and other genes involved in immune signaling (Fig. 3B). Notably, we observed induction of genes involved in DDX58/IFIH1 (RIG-I/MDA5)-mediated induction of IFNa/b at 4h, followed by induction of negative regulators of this pathway at 24h (Fig. 3B). Similarly, an enrichment for genes involved in response to cytokine stimuli was detected at 4h but not at 24h post treatment. These observations of a strong early response followed by a decline by 24h upon IFN stimulation are consistent with observations in the black flying fox (*Pteropus alecto*) [9].

**Fig. 3.**
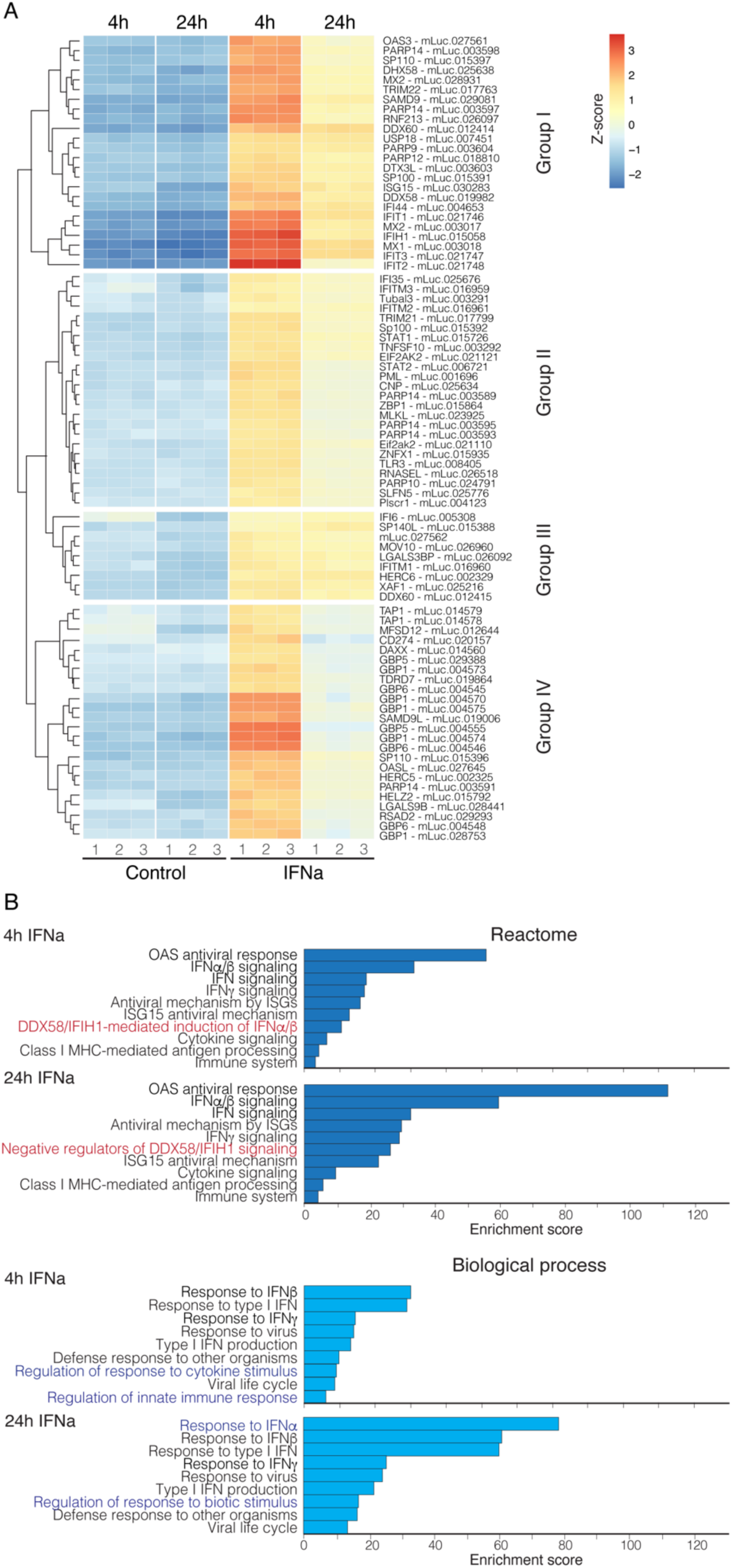
Gene expression profiles and enrichment analyses. **A**) Heatmap shows the 140 genes with highest expression variance across samples in vsd transformed data (clustering method: “euclidean”). Embryonic fibroblast cells presented four different profiles of gene induction dynamics through time: i) genes with rapid induction and decline at 24h (Group I); ii) genes with mild induction at 4h and low decline at 24h or with stable induction (Groups II and III); and iii) genes with rapid induction at 4h and rapid decline (Group IV). **B**) Bar graphs show the results of functional overrepresentation analysis (ORA) on differentially expressed genes at 4h and 24h post IFNa treatment. Although most terms are shared between the two time points, we found the DDX58/IFIH1 pathway to be active at 4h post IFNa stimulation, whereas it is subject to negative regulation at 24h. Similarly, the response to cytokine stimuli is present at 4h post treatment, but not at 24h.

### TE contribution to ISG transcript structure

To improve detection of potentially unannotated IFN-induced TE-derived transcripts and isoforms, we conducted genome-guided transcriptome reassembly on our combined RNA-seq dataset using StringTie2 [43] (Supplementary Data 1), and annotated assembled genes based on homology using the SwissProt database. This yielded an expanded transcriptome with 68,110 transcripts (32,137 transcripts matching an annotated gene). After performing pairwise differential expression analyses (Table S3) we identified 1243 IFN-inducible transcripts corresponding to 836 StringTie genes (740 IFN-inducible transcripts corresponding to 449 StringTie genes matching Swissprot) at 4h, and 717 transcripts corresponding to 500 StringTie genes (385 transcripts, 239 matching Swissprot) at 24h. Of these, 606 transcripts corresponding to 392 genes (358 transcripts, 214 genes matching Swissprot) were shared between 4h and 24h treatment time.

We used our improved transcriptome reassembly to investigate the contribution of TEs to both constitutive and IFN-inducible transcript structures. First, we identified transcripts that contained exonized TEs based on the overlap of exon (>50% sequence length) and annotated TE features. Considering all expressed (TPM ≥ 0.5) multi-exon transcripts, we found a subset of 1039 transcripts corresponding to 749 StringTie genes (648 transcripts and 470 genes with homology to annotated genes in the SwissProt database) that contained at least one TE-derived exon (Table S4).

Focusing first on transcripts reconstructed from constitutively expressed genes, we identified *EEF1A1* as an example where 271 bp (representing part of the third and fourth exons; coding exons 2 and 3) is annotated as derived from a Zisupton DNA transposon. Since Zisupton DNA elements have likely been lost during the tetrapoda radiation [44], and their presence and activity has been confirmed only in fish species among vertebrates, we further verified this finding through multiple BLAST searches (Supplementary Data 2). First, we aligned the StingTie transcript (STRG.21546.2, Scaffold 10: 12654609-12761039) to the transcript sequence deposited on the NCBI database (XM_006089332.3). After confirming the match between the two transcripts, we used the deposited mRNA sequence as a query against the Repbase transposable element database [45] and against bats and mammalian mRNA collections. All *Microchiroptera*, *Vespertilionidae* in particular, share highly homologous coding exons 3 and 4 (Supplementary Data 3), suggesting that the exonization event occurred before or during the *Microchiroptera* and *Pteropodinae* radiation. Despite the shared homology, no Zisupton matching features were found in the human or other euarchonta *EEF1A1* homologues. Therefore, our analysis successfully identified bat *EEF1A1* as an example of a gene that has likely been altered by a bat-specific TE.

We next filtered for IFN-inducible transcripts containing TE-derived exon(s), and found 44 transcripts from 34 genes (16 of which annotated by SwissProt homology) induced at 4h, and 31 transcripts from 24 genes (11 annotated) induced at 24h (Table S4). These included genes with established roles in immune responses, like *PARP9*, *SLFN5* (Fig. S2A) and a candidate novel isoform of *IFITM2* that has not been previously identified (Fig. S2B). This analysis reveals that numerous constitutively expressed and IFN-inducible genes have acquired exons from bat-specific TEs either in coding regions or UTRs (Table 1).

**Table 1.**
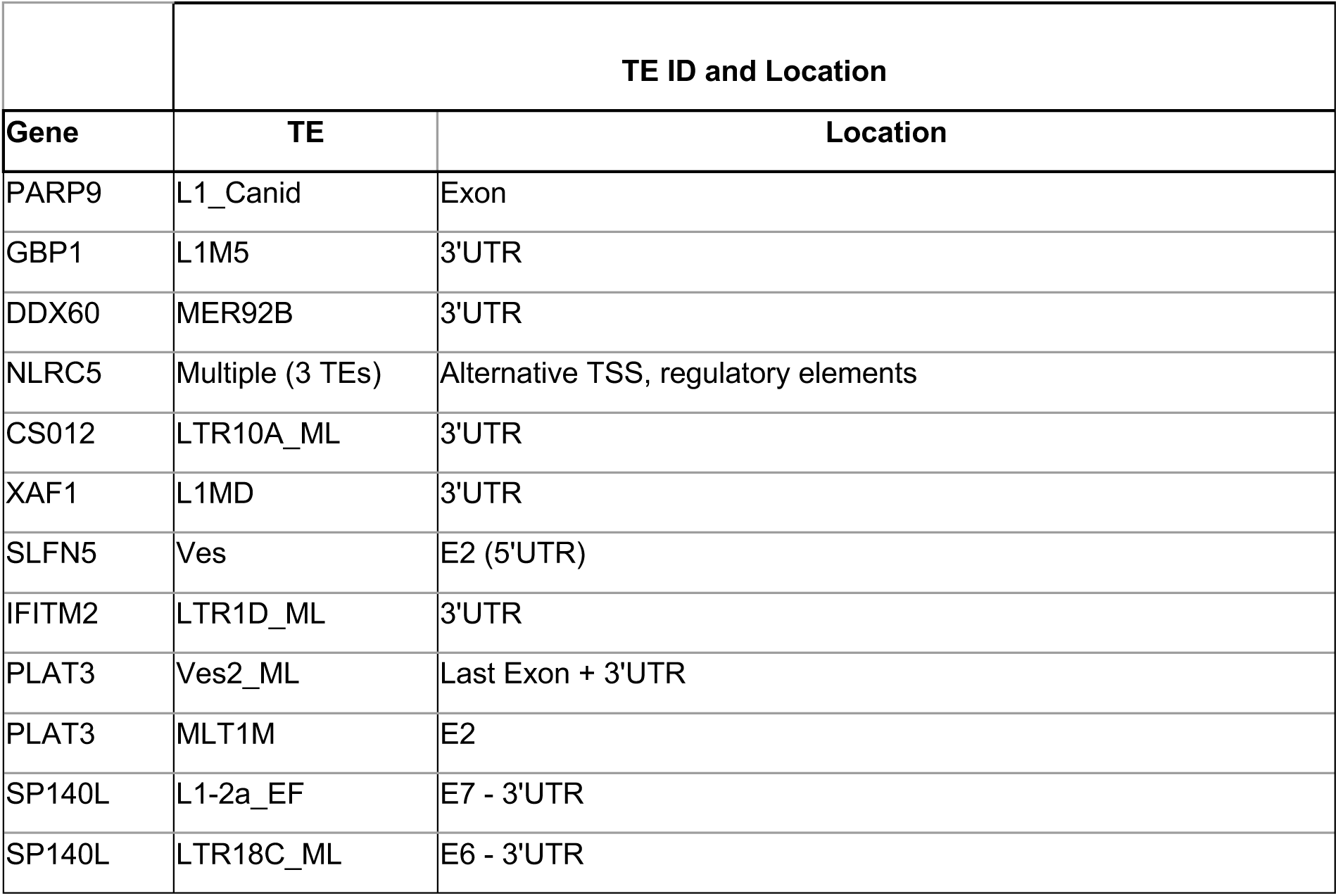
List of candidate genes with TE-derived exons. Transposable element (TE) exonization events were predicted by intersecting coordinates of annotated TEs and de-novo transcriptome assembly of RNA-seq data for *Myotis lucifugus*. Each candidate was inspected in our custom UCSC genome browser track (https://genome.ucsc.edu/s/GiuliaPasquesi/myoLuc2_HiC) and selected if it showed multiple lines of support (RNA expression, chromatin profile, additional gene/isoform annotation).

|  | TE ID and Location |  |
| --- | --- | --- |
| Gene | TE | Location |
| PARP9 | L1_Canid | Exon |
| GBP1 | L1M5 | 3'UTR |
| DDX60 | MER92B | 3'UTR |
| NLRC5 | Multiple (3 TEs) | Alternative TSS, regulatory elements |
| CS012 | LTR10A_ML | 3'UTR |
| XAF1 | L1MD | 3'UTR |
| SLFN5 | Ves | E2 (5'UTR) |
| IFITM2 | LTR1D_ML | 3'UTR |
| PLAT3 | Ves2_ML | Last Exon + 3'UTR |
| PLAT3 | MLT1M | E2 |
| SP140L | L1-2a_EF | E7 - 3'UTR |
| SP140L | LTR18C_ML | E6 - 3'UTR |

We next aimed to identify potential examples of co-opted TE-derived proteins (such as syncytins; [46]) which may not be masked by RepeatMasker due to their age. We used *tblastx* to query expressed StringTie transcripts against reference TE protein sequence libraries specific for retrotransposons and DNA elements (see Methods). We found a total of 7 StringTie genes with a known annotated ortholog based on the concatenated StringTie transcriptome assembly, and 156 StringTie genes based on intact ORF prediction (Table S5; Methods). By using the intact ORF prediction approach we identified 9 ISGs (i.e., *GBP1*, *DDX58* and *PARP14*) with at least one retrotransposon-derived feature, mostly from ERVK and L1 LINEs. Most of these TE-derived sequences reside in the last exon, where they provide both the stop codon and the 3’UTR, or novel coding and/or regulatory sequences. Finally we followed the same approach to identify novel protein-coding sequences matching known viral proteins that could represent domesticated viral proteins [47]. By leveraging the gEVE and a custom syncytin protein database we found a total of 3 constitutively expressed StringTie genes with homology to syncytin proteins and 9 with homology to either a pol, gag, retrotransposase or AP viral proteins (Table S6). Only one pol (RVT1)-derived gene, *UBP18*, was differentially expressed upon IFN treatment .

In addition to examining the TE contribution to protein-coding sequences, we also searched for examples of TE-derived promoters. We collapsed StringTie transcript coordinates to their transcription start site (TSS), and intersected transcript TSSs with the generated TE annotation. We identified a total of 11 transcripts with a TSS deriving from a TE that are IFN-inducible at 4h, 9 of which are shared with the 24h subset (Table S7). Most of these belong to genes known to be involved in immune function and regulation, like *NLRC5* (Fig. 4), *EIF2AK2*, *GBP1*, *MX1*, *MX2*, *PHF11*, *SAMD9* and *XAF1*, while others like *PARP14* and *CS012* are not canonical ISGs. Notably, we also observed recurrent usage of TE-derived promoters for histocompatibility loci located in Scaffolds 13 and 20, where 78 transcripts (corresponding to 21 annotated genes) at 4h and 7 transcripts (corresponding to 3 annotated genes) at 24h have TE-derived TSSs or coding exons. While the histocompatibility locus may be prone to sequence assembly artifacts related to its highly polymorphic and complex structure, our analysis suggests that TEs may have influenced the evolution of the bat histocompatibility locus, which has been proposed to underlie bat-specific immune function [48, 49].

**Fig. 4.**
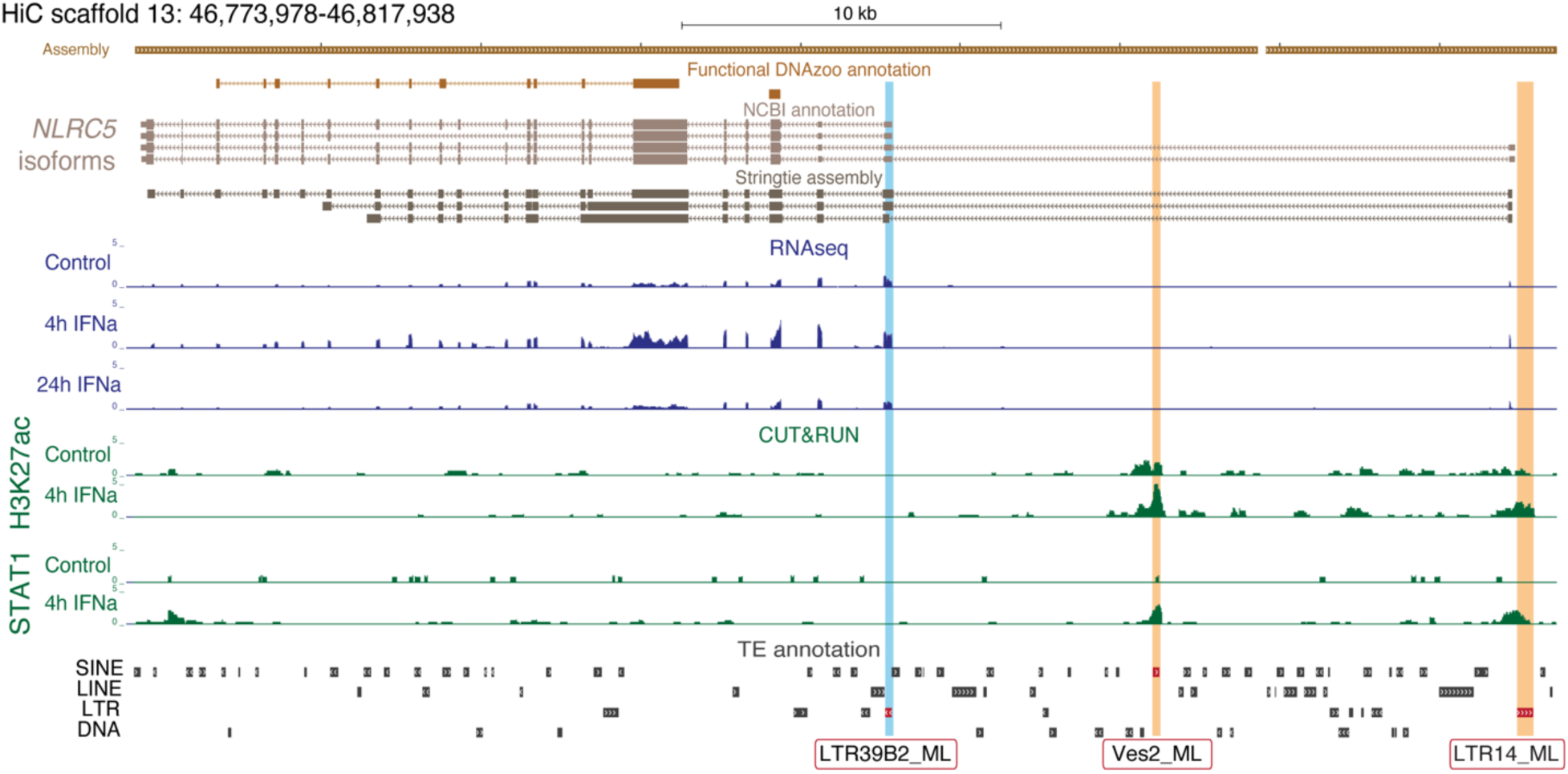
TE exonization events in *Myotis* ISGs. Custom UCSC genome browser screenshot of the NLRC5 locus, where one exon (light blue highlight) represents a potential alternative transcription start site (TSS) deriving from a Myotis-specific LTR39B2_ML retrotransposons. RNAseq coverage at the promoter region suggests upregulation of the transcript at 4h post IFNa treatment, consistent with its role in immune responses, and lower expression in 24h post treatment samples compared to unstimulated cells. We also identified 2 potential TE-derived regulatory elements (orange highlight) in intronic or upstream regions of the NLRC5 locus that show increase in H3K27ac and STAT1 CUT&RUN signal at 4h post IFNa treatment.

### Epigenetic profiling upon IFN stimulation

Having analyzed the contribution of TEs to ISGs, we next asked how TEs contribute to inducible regulatory elements defined by H3K27ac, POLR2A, or STAT1 activity in response to IFN. We observed an increase in STAT1 signal at the predicted promoters for *IRF9* and *PSME2* in response to type I IFN (Fig. 5A). Using spike-in normalized CUT&RUN data at 0 and 4h, we used DESeq2 to define IFN-inducible regulatory elements, as performed previously [22]. Unexpectedly, we did not observe robust IFN-inducible chromatin changes that are characteristic of IFN-stimulated cells from other species [22, 50]. We did not identify any H3K27ac inducible elements with an FDR < 0.05 (Fig. 5B; Table S8), and instead defined a set of 1113 elements showing an increase of H3K27ac signal with a relaxed significance threshold of unadjusted *p*-val < 0.1 (Fig. S3A). This set of IFN-inducible elements was enriched for interferon-stimulated response element (ISRE) motifs (E-value 1.43×10^-10^) (Table S9), consistent with their activation by IFN stimulation. Thus, while our CUT&RUN analysis successfully identified some elements showing IFN-inducible activity, our analysis reveals surprisingly modest chromatin-level changes, despite robust ISG induction according to RT-qPCR from RNA taken from the same fraction of cells (Fig. S1).

**Fig. 5.**
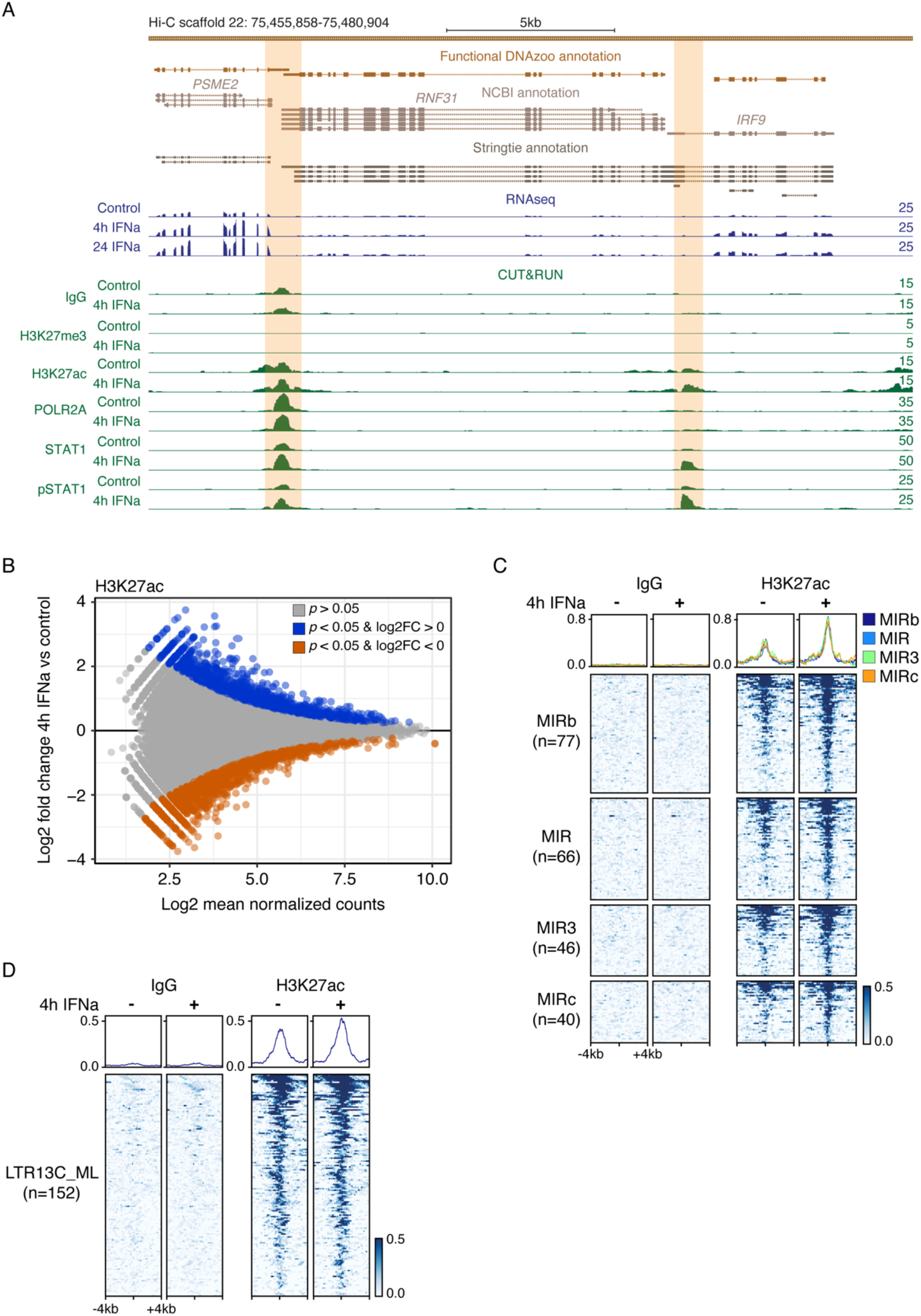
Epigenomic profiling of untreated and IFNa-stimulated embryonic fibroblast primary cells. **A)** Genome browser view of the *IRF9* and *PSME2* loci. RNA-seq and CUT&RUN tracks are normalized per million reads. Signal track maxima are indicated on the right of each track. IFNa-inducible (*p*-val < 0.10, log2FC > 0) STAT1 peaks are highlighted orange. **B)** MA plot of IFNa-inducible (unadjusted *p*-val < 0.10, log2FC > 0, blue) and IFNa-repressed (unadjusted *p*-val < 0.10, log2FC < 0, orange) H3K27ac regions from H3K27ac CUT&RUN. Regions with an unadjusted *p*-val > 0.10 are shown in grey. Log2 fold change values were shrunken using the apeglm function v1.8.0 [83] .**C)** Heatmaps showing normalized CUT&RUN signal (signal per million reads) over IFNa-inducible (unadjusted *p*-val < 0.10), H3K27ac-marked MIR families. **D)** Heatmaps showing normalized CUT&RUN signal (signal per million reads) over H3K27ac-marked LTR13C_ML families.

Of these regions, we found that 466 out of 1113 fully overlapped at least one TE (Table S10). Additionally, we identified 766 inducible, STAT1-bound TEs that fall within 100kb of an ISG (Table S11). This includes an LTR14_ML element that may be functioning as the promoter for the *NLRC5* locus in addition to an intronic Ves2_ML SINE element (Fig. 4). However, in contrast to previous studies in other species [22–26], we did not observe any overrepresented TE families within this set (Fig. S3B; Table S12). The only subfamilies that overlapped more than 10 IFN-inducible H3K27ac regions correspond to the ancient mammalian MIR and L2 families that predate the evolution of bats (Fig. 5C; Fig. S3C). Querying H3K27ac regions from either untreated or IFN-stimulated conditions independently, we observed only very modest enrichment of the MIR3, AmnSINE1, and LTR13C_ML families (Fig. 5D; Fig. S4A, S4B). Taken together, our analysis indicates that TEs contribute to hundreds of regulatory elements involved in IFN signaling, but in contrast to studies in other species, we did not identify enrichment of lineage-specific TE families within IFN-inducible regulatory elements. However, given that our CUT&RUN analysis revealed a relatively minimal set of inducible regulatory elements at a genome-wide level, we were limited in our ability to identify enriched TE families.

## DISCUSSION

Our study characterizes the transcriptional and epigenetic dynamics of bat TEs in the IFN response in *M. lucifugus* cells. To facilitate our ability to map TEs in our functional genomic data, we conducted both RNA-seq and CUT&RUN using 150bp paired end reads, and generated an improved repeat annotation using a chromosome-scale assembly. Our analyses revealed that TEs have shaped the IFN-inducible transcriptome, but we did not find strong evidence for a global role for TEs in shaping associated epigenetic changes. Functional studies will be necessary to validate whether any of the elements identified in this study have significance for bat immunity, but given the growing number of validated examples in human and mouse, it is likely that some TEs have been co-opted for innate immune function in bats.

For our study, we generated a matched transcriptomic and epigenomic datasets profiling the type I IFN response in *M. lucifugus* primary cells. Our transcriptomic analysis of the IFN response in *M. lucifugus* embryonic fibroblasts confirms previously reported features of bat innate immunity. We found that these cells respond to IFN stimulation at the transcriptional level, with a stronger and broad induction of ISGs at earlier time points (4 hours post treatment). We also found that only a small subset of genes that were overexpressed at 4h maintain high expression levels at 24h, whereas most genes show a reduction in expression to lower levels or levels similar to those recorded in unstimulated cells. This is in agreement with gene expression profiling in *Pteropus alecto* [9]. In parallel to gene expression, we characterized expression profiles of TE-derived transcripts, and found similar trends. Total TE expression was higher at 4h post IFN treatment, with more and more diverse TE families being differentially expressed both in comparison to unstimulated and 24h cells. While the expression of these transcripts is not directly indicative of function, IFN-inducible expression of bat-specific TE families may act as a source of non-coding transcripts that can further activate innate immune pathways, akin to the “viral mimicry” pathway characterized in human cancers [19, 20].

We also explored whether specific TEs may have affected the transcript structure of host genes, by screening for gene transcripts that share homology with transposable elements or viral proteins in coding regions and transcription start sites (TSS). Our genome-guided transcriptome assembly identified multiple instances of TE-derived and viral protein-derived exonization events, both as alternative (*IFITM2*) or conserved (*PARP9*) exons and TSS (*NLRC5*). Some of these transcripts represent canonical (*PARP9*, *DDX60*) or non-canonical (*PLAAT3*, *SP140L*) ISGs. These analyses provide strong evidence that TEs have been co-opted into the exons of bat ISGs, and some of these exonization events may have significant functional consequences. For example, our analysis identified multiple *Myotis*-specific TE insertion and exonization events affecting the *NLRC5* gene. NLRC5 has been identified as a key regulator of MHC class I-dependent immune responses [51], and may be involved in the regulation of inflammasome activation and type I IFN responses [52]. Further studies are needed to validate the potential effects of these TE-derived sequences, but it is possible that *Myotis*-specific TEs have altered NLRC5 function and/or regulation.

We also identified instances of TE-derived constitutively expressed genes. We verified through multiple BLAST and sequence alignments that the ∼100 amino acids of the EEF1A1 protein of *Microchiroptera* and likely *Pteropodinae* bats derived from the exonization of a Zisupton DNA transposons. Although Zisupton DNA transposons are not abundant in the *M. lucifugus* genome (we recovered only one family present, with 174 copies), it is possible that they were horizontally transferred into bat genomes as has happened in multiple fish species [44]. Age analysis of Zisupton genomic copies suggests that they were likely recently introduced (divergence from consensus ∼18), but were able to expand only for a short period of time (peak of divergence from the consensus sequence at 4-9), when other DNA elements have been active. This example uncovered by our analysis highlights the possibility that TEs have shaped other aspects of bat biology in addition to genes involved in immune function.

While RNA-seq profiling has been applied by an increasing number of studies to profile bat immunity at the transcriptomic level, no study to date has characterized bat immunity at the epigenomic level. Unexpectedly, while our RNA-seq analysis of *M. lucifugus* cells coincided with strong transcriptional response to IFN treatment, we observed relatively modest chromatin changes based on CUT&RUN epigenomic profiling. This observation contrasts with robust chromatin changes typically observed in IFN-treated cells from other mammalian species [22,53,54]. As a result, we did not observe strong evidence supporting family-level TE regulatory induction as observed previously in other species such as human [50] and cow [22], partly due to the lack of clearly inducible elements at a genome-wide level.

There are multiple potential explanations for our observation of a relatively modest IFN-inducible epigenetic response in bat cells. First, our study is one of the first to conduct CUT&RUN on bat cells, and the antibodies used in this study have not been fully validated in *M. lucifugus*. However, while the antibody used for STAT1 may exhibit poor recognition of bat ortholog, histones and POLR2A are highly conserved and expected to be targeted effectively by standard antibodies. Second, our study involved stimulating derived embryonic fibroblast cell culture with recombinant universal type I IFN. While these conditions nonetheless showed strong transcriptional induction of ISGs, it is possible that chromatin dynamics are different during endogenous activation of IFN responses *in vivo*.

Finally, our observations may reflect a unique attribute of bat immunity, consistent with the idea that some bat species exhibit constitutive IFN expression [7, 55]. Although the type I IFN locus of *M. lucifugus* is poorly characterized, we were able to annotate at least 11 uncharacterized genes that likely reflect the expansion by gene duplication of the IFNw cluster (Fig. S5) [56], whereas no IFNa genes were identified. Of the 11 IFNw paralogues, 5 showed evidence of constitutive low expression in unstimulated cells, and induction at 4h post treatment (Fig. S5). These observations suggest a scenario where *Myotis* epigenomes are “primed” due to constitutive expression of IFN, and may be capable of driving robust transcriptional activation without exhibiting epigenetic changes typically associated with inducible chromatin activity, such as increased H3K27ac or POLR2A levels.

## CONCLUSIONS

Our study provides a first systematic investigation of the contribution by TEs to the bat type I IFN response. We uncover numerous examples of TE-derived transcripts, alternative exons, and regulatory elements that shape the genomic response to IFN in *M. lucifugus*. Our study suggests that TEs in other bat lineages such as *Pteropus* and *Rhinolophus* may also shape IFN-inducible transcriptomes, which may motivate functional studies to determine their biological significance in the context of bat immunity. Our findings lend additional support for a widespread role for TE co-option in shaping the evolution of species-specific immune responses.

## METHODS

### Transposable Element identification and analysis

*Myotis lucifugus* genomic repeat elements were annotated according to homology-based and *de novo* identification approaches. Although repeat elements have been extensively annotated in bat species including *Myotis lucifugus*, we performed *de novo* TE identification using RepeatModeler2 (included in dfam-tetools-1.1) [37] and HelitronScanner v1.1 [38] to match the newly released, highly contiguous chromosome-level assembly of the little brown bat genome (myoLuc2_HiC) [36,57,58]. Given their highly repetitive nature, TE loci are hard to assemble and often incomplete; therefore genome assemblies that rely on long read assembly strategies (i.e., HiC) [36, 59] are better suited for capturing full length elements over contiguous chromosome-level scaffolds.

Briefly, we performed de novo TE identification using RepeatModeler and HelitronScanner, then combined the two libraries as a single little brown bat *de novo* library that was used for homology-based TE annotation using RepeatMasker v4.1.0 [40]. To maximize element identification we followed a custom multi-step mapping strategy [60] using multiple libraries as reference for the masking process in the following order: (i) bat specific repeats included in the Repbase library provided with RepeatMasker; (ii) a bat specific library provided by Dr. Cosby; (iii) our *de novo* little brown bat library; (iv) the entire tetrapoda Repbase library provided with RepeatMasker [45].

### Cell lines and treatment

*Myotis lucifugus* primary embryonic fibroblast cells were a gift from Mario Capecchi. Cells were grown at 37°C and 5% CO­and passaged in DMEM (ThermoFisher #10566016) supplemented with 10% FBS, 5% MEM nonessential amino acids, 100 U/mL Penicillin-Streptomycin, and 1 mM sodium pyruvate. Cells were seeded into six-well plates at an optimized density of 2×10^5^ cells per well in 2ml of culturing media (or 2×10^6^ cells per 15 cm dish for CUT&RUN). The following day (or 48h for CUT&RUN) cells were treated with 1000U/ml of Universal Type I IFNa resuspended in DPBS (PBL Assay Science #11200) in 2ml of culturing media; control cells were treated with equivalent volume of DPBS in 2ml of media. At four hours and 24 hours post treatment cells were harvested for RNA extraction (four hours for CUT&RUN).

### RNA isolation and library preparation for RNA-seq

Following media removal, cells were washed with 1ml of DPBS and detached by adding 400ul of 0.25% trypsin per well. Following a 10 minutes incubation at 37°C, trypsin was neutralized with 1.6ml of culturing media. Cell suspensions were transferred into 1.7ml tubes and pelleted by centrifugation at 300xg for 5 minutes. Cells were then lysed in 300ul of RNA lysis buffer (Zymo Research #R1060-1-50), and stored at -80°C until RNA extraction was performed using the Quick-RNA MiniPrep kit (Zymo Research #R1054), following manufacturer’s instructions.

Total RNA samples for each time point and condition were prepared in three biological replicates as described above. A NanoDrop One spectrophotometer (Thermo Fisher Scientific) was used to determine RNA concentration and quality; all samples passed quality assessment. PolyA enrichment and library preparation was performed using the KAPA mRNA HyperPrep Kit (Kapa Biosystems #8098115702) according to the manufacturer’s protocols. Briefly, 500 ng of RNA was used as input, and single-index adapters (Kapa Biosystems #08005699001) were added at a final concentration of 10 nM. Purified, adapter-ligated library was amplified for a total of 11 cycles following the manufacturer’s protocol. The final libraries were pooled and sequenced on an Illumina NovaSeq 6000 (University of Colorado Genomics Core) as 150bp paired-end reads.

### CUT&RUN sample and library preparation

CUT&RUN pulldowns were generated using a protocol from [61, 62]. All buffers were prepared according to the “High Ca^2+^/Low Salt” section of the protocol using 0.04% digitonin (EMD Millipore #300410). 5×10^5^ viable cells were used for each pulldown. The following antibodies were used: rabbit anti-mouse IgG (1:100, Abcam #ab46540), rabbit anti-H3K27me3 (1:100, Cell Signaling #9733), rabbit anti-H3K27ac (1:100, Millipore #MABE647), rabbit anti-pRPB1-Ser5 (1:50, Cell Signaling #135235S), rabbit anti-STAT1 (1:100, Cohesion #3322), rabbit anti- pSTAT1-Ser727 (1:100, Active Motif #39634). pAG-MNase (prepared as in [61, 62]) was added to each sample following primary antibody incubation at a final concentration of 700 ng/mL. Chromatin digestion, release, and extraction was carried out according to [61, 62]. Yeast spike-in DNA (gift from Steven Henikoff) was added to the quenching (“1× STOP”) buffer for a final concentration of 100 pg/mL. Pulldown success was determined by Qubit dsDNA High Sensitivity and TapeStation 4200 HSD5000 before proceeding with library preparation.

Libraries were generated using a modified protocol for use with the KAPA HyperPrep Kit. Briefly, the full volume of each pulldown (50 uL) was used to generate libraries according to the manufacturer’s protocol with the following modifications. Freshly diluted 0.200 uM single-index adapters (Kapa Biosystems #08005699001) were added to each library at a low concentration (9 nM) to minimize adapter dimer formation. Adapter-ligated libraries underwent a double-sided 0.8X/1.0X cleanup with KAPA Pure Beads (Kapa Biosystems #07983280001). Purified, adapter-ligated libraries were amplified using the following PCR cycling conditions: 45 s at 98°C, 15x(15 s at 98°C, 10 s at 60°C), 60 s at 72°C. Amplified libraries underwent a double-sided 0.8X/1.0X cleanup. The final libraries were quantified using Qubit dsDNA High Sensitivity and TapeStation 4200 HSD5000. Libraries were pooled and sequenced on an Illumina NovaSeq 6000 (Novogene) as 150bp paired-end reads.

### Paired RT-qPCR

5×10^5^ viable cells from the same CUT&RUN populations (untreated and 4h IFN) were used to extract RNA for RT-qPCR analysis to confirm induction of IFN-inducible genes prior to CUT&RUN library preparation. Cells were lysed in 300ul of RNA lysis buffer (Zymo Research, #R1060-1-50). Prepared lysates were stored at -80°C until RNA extraction was performed using the Quick-RNA MiniPrep kit (Zymo Research #R1054), following the manufacturer’s instructions.

Total RNA samples for each time point and condition were prepared in three biological replicates as described above. A NanoDrop One spectrophotometer (Thermo Fisher Scientific) was used to determine RNA concentration and quality; all samples passed quality assessment. RNA expression levels for *CTCF*, *STAT1*, and *IFIH1* were quantified using the Luna Universal One-Step RT-qPCR Kit (New England Biolabs #E3005L) according to the manufacturer’s instructions. In brief, for each reaction 25 ng of RNA was combined with 5ul 2× Luna Universal One-Step Reaction Mix, 0.5ul 20× Luna WarmStart RT Enzyme Mix, 0.4ul 10uM forward primer, and 0.4ul 10uM reverse primer. Reactions were amplified using a CFX384 Touch Real-Time PCR Detection System (Bio-Rad) with the following PCR cycling conditions: 10 min at 55°C, 1 min at 95°C, 40x(10 s at 95°C, 30 s at 60°C). On-target amplification was assessed by melt curve analysis. Two biological replicates were included for each treatment condition, and each biological replicate was run in technical duplicate. Statistical significance was assessed using a two-tailed paired Student’s t-test with a threshold of *p*-val < 0.05.

### Transcriptome analyses

Paired-end 150bp read length FASTQ files were quality and adapter trimmed using BBDuk v.38.05 [63]; quality check was performed using FastQC v0.11.8 [64] and inspected through MultiQC v1.7 [65]. Filtered FASTQ files were then mapped to the myoLuc2_HiC genome using a 2-pass approach in STAR v2.7.3a [66]. STAR was run following default parameters and allowing for multi-mapping reads (with options ‘*–outAnchorMultimapNmax 100 – winAnchorMultimapNmax 100 –outFilterMultimapNmax 100*’), a requisite for the inclusion of TE-mapping reads in the output files. The annotation file available on DNAzoo, here referred to as “functional annotation” (www.dropbox.com/sh/xt300ht42mihjov/AADoENW7RTvR3jTh1a8qUOmRa) was used as reference for the mapping process. For the second pass of mapping we filtered out novel junctions that mapped to the mitochondrial genome, HiC_scaffold_93 based on the most likely alignment hit to the reference mitochondrial genome performed using LASTZ v1.02.00 [67]. Resulting alignment files in sorted .*bam* format were then provided as input for TE and gene expression quantification in TETranscripts v2.1.4 [42] using the same gene annotation and our custom TE annotation derived from RepeatMasker. Pairwise differential expression analyses at 4h and 24h post IFN treatment were performed in DESeq2 v1.32 [68]. Functional enrichment analyses of differentially expressed genes (adj. *p*-val < 0.05, log2FC > 1.5) were performed using the WebGestalt web tool [69].

### Genome guided transcriptome assembly and analysis

Short-read RNA-seq alignment files generated by running STAR (see previous paragraph) were merged, sorted and indexed using SAMtools v1.10 [70], and the resulting .*bam* file was used as input for genome guided transcriptome assembly in StringTie v1.3.3b [43]. StringTie was run following default parameters, except that the minimum number of spliced reads required to align across a junction was increased from 1 to 5 using option ‘*-j 5*’. The resulting StringTie gtf output (Supplementary Data 1) was then converted to FASTA format using the *gffread* utility using options ‘*-M, -F, and -Z*’ included in Cufflinks v.2.2.1 [71]. We followed an homology-based approach to annotate assembled StringTie genes and isoforms. StringTie gene sequences were queried against the SwissProt database [72] through the *blastx* search algorithm in BLAST 2.12.0+ [73] using options ‘*-max_target_seqs 1 -evalue 10*‘. Matches were then filtered for shared sequence identity equal to or greater than 50%.

To find transcripts with coding regions that intersect TEs, we applied BEDTools v2.28.0 [74] to filter for events where at least 50% of the sequence of one exon derived from a TE with option ‘*- f 0.5*’. Briefly, exon coordinates were extracted from the StringTie annotation and intersected with our custom TE annotation for *Myotis lucifugus*. To narrow down the list of candidate StringTie transcripts and limit redundant of false positive matches, the output was filtered for multi-exon transcripts with a transcripts per million (TPM) value equal to or higher than 0.5. TPM values were quantified at the isoform level by RSEM v1.3.0 [75]. Filtered StringTie transcript candidates were cross-referenced against the results of differential expression analysis in DESeq2 v1.32 [68] (Table S3) to identify TE exonization events in ISGs. Candidates ISGs were verified by visually inspecting candidates against the DNAzoo and NCBI annotations for *Myotis lucifugus* and RNA-seq coverage tracks. To identify TEs that might be contributing to alternative transcriptional start sites (TSS), we used a custom python script to extract the TSS coordinates from the StringTie annotation and intersected this collapsed file with the TE annotation file as previously described.

Finally, we queried StringTie-derived assembled transcripts against databases of DNA transposons and retrotransposons extracted from the Repbase repository of TE reference sequences [45]. *tblastx* was used according to previously specified parameters and resulting hits were further filtered for alignments greater than or equal to 300bp with a sequence identity greater than or equal to 90%. To identify any transcripts with intact protein-coding sequences of viral origin, we ran *blastx* against i) the gEVE repository of retroviral proteins [76] and ii) a custom database of syncytin proteins collected from the ncbi repository [77]. The output of gEVE blast was filtered for alignments greater than or equal to 200bp with a shared sequence identity greater than or equal to 50%. The output of syncytin blast was filtered for alignments greater than 100bp with shared sequence identity of 50% and above. The same blast analysis for TE sequences and protein databases was carried out on identified open reading frames (ORFs) larger than 50aa found by running the function *usearch* [78] (*usearch -fastx_findorfs -orfstyle 7 -mincodons 16)* on the StringTie assembled transcripts file.

### CUT&RUN analysis

Adapters and low quality reads were trimmed using BBDuk v38.05 [63] using options *‘ktrim=r k=34 mink=11 hdist=1 tpe tbo qtrim=r trimq=10*’. Trimmed reads were aligned to the myoLuc2_HiC assembly using Bowtie 2 v2.2.9 [79] with options ‘*--local –very-sensitive-local – no-unal –no-mixed –no-discordant -I 10 -X 700*’, and only uniquely mapping reads with a minimum MAPQ of 10 were retained. Fragments aligning to the mitochondrial genome were removed. Trimmed reads independently aligned to the *S. cerevisiae* assembly (GCF_000146045.2) using Bowtie 2 v2.2.9 [79] with options *‘--local –very-sensitive-local –no-unal –no-mixed –no-discordant –no-overlap –no-dovetail -I 10 -X 700*’. myoLuc2_HiC read depth was normalized according to the number of fragments aligned to the *S. cerevisiae* assembly for each sample, and normalized bigWigs corresponding to read coverage per 1 million normalized reads were generated using deepTools v3.0.1 [80, 81] for heatmap visualization.

Peak calling was performed using complete and size subsetted alignment files with MACS2 v2.1.1 [82] in a two-step process where separate sets of peaks were called with 1) single-end options ‘*--format BAM --shift=-75 --extsize=150*’ and 2) paired-end option ‘*--format BAMPE*’. For both modes only peaks with an unadjusted *p*-val < 0.01 were retained. Peaks from each mode were subsequently merged. IgG peaks were subtracted from each pulldown peak set to minimize background. Only the top 20,000 peaks by descending MACS2 peak score were retained for further analysis.

To identify IFN-inducible CUT&RUN peaks, the top 20,000 peaks for all samples for a particular pulldown (across all replicates, untreated and IFN-stimulated) were concatenated into a single list, and aligned fragments from each individual sample were counted for all peaks using BEDTools v2.28.0 [74]. IFN-inducible peaks were called using DESeq2 v1.26.0 [68], however we were unable to identify peaks that were significantly upregulated in response to IFN with an FDR < 0.10. We therefore took a more relaxed approach, retaining all peaks with an unadjusted *p*-val < 0.10 and log2FC > 0. log2FC values were shrunken using the apeglm function v1.8.0 [83] for visualization. Motif analysis was performed using XSTREME v5.4.1 [84] with options *‘-- minw 6 --maxw 20 --streme-motifs 20 --align center*’ querying against the JASPAR CORE 2018 vertebrates database [85].

To assess the contributions of TEs in regulating the IFN response, we intersected IFN-inducible H3K27ac peaks with all annotated TEs, requiring that all reported TEs are fully contained within the peak. We further characterized the regulatory landscape by identifying STAT1-marked TEs as a function of distance to the nearest ISG (FDR<0.05, log2FC>1.5) transcriptional start site using BEDTools v2.28.0 [74]. To assess family-level enrichment, GIGGLE v0.6.3 [86] was used to create a database of TEs in the myoLuc2_HiC genome using the custom TE database as described above. IFN-inducible H3K27ac peaks were then queried against the TE database. Results were ranked by descending Giggle enrichment score, and enriched TE families were identified according to the odds ratio, Fisher’s two-tailed *p*-val, and number of overlaps. The TE heatmaps were prepared by selecting elements within various families that overlapped either IFN-inducible H3K27ac regions or any H3K27ac regions from untreated or IFN conditions. Signal from *S. cerevisiae* spike-in, CPM normalized bigwigs was plotted as heatmaps using deepTools v3.0.1 [80, 81].

## DECLARATIONS

### Ethics approval and consent to participate

Not applicable.

### Consent for publication

Not applicable.

### Availability of data and materials

Newly generated RNAseq and CUT&RUN raw files have been deposited under the GEO SuperSeries accession GSE200833. Processed data, including TE annotation, can be visualized as a UCSC genome browser custom track here: https://genome.ucsc.edu/s/GiuliaPasquesi/myoLuc2_HiC.

### Authors’ contributions

EC and GP designed the study. GP and CK performed experiments. GP, CK, AO and EC analyzed data and interpreted results. GP and CK created and edited figures and tables. EC, GP and CK wrote the manuscript with input from all co-authors. All authors gave final approval for publication.

## Acknowledgements

Unpublished genome assemblies are used with permission from the DNA Zoo Consortium (dnazoo.org). *Myotis lucifugus* primary embryonic fibroblast cells were a kind gift of Mario Capecchi (University of Utah). We thank the BioFrontiers Computing core for technical support during this study.

