## Supplementary Figures for "Transcriptional dynamics of transposable elements in the type I IFN response in *Myotis lucifugus* cells"

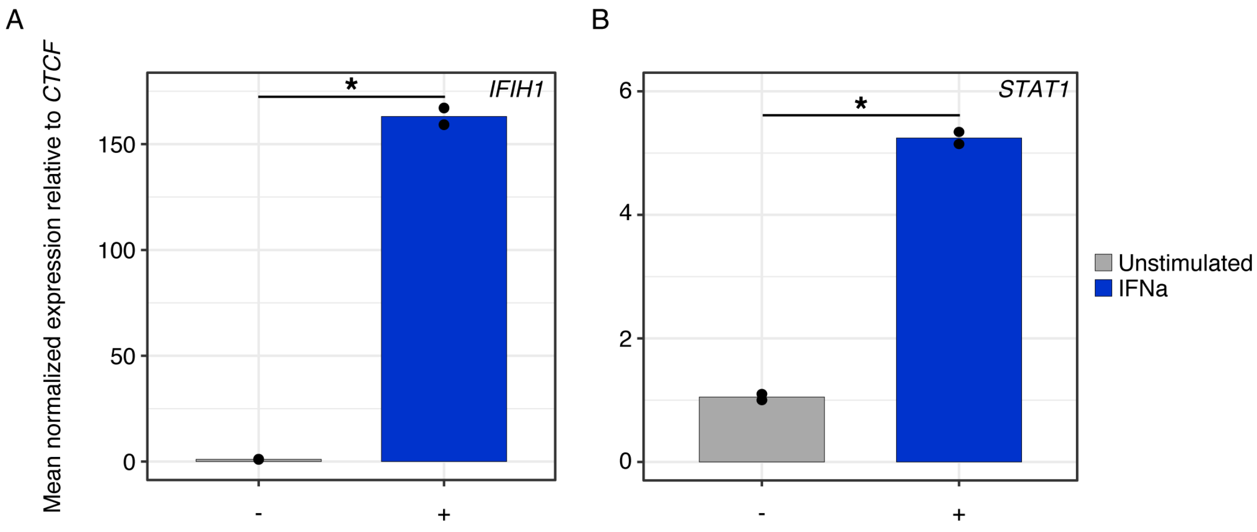


**Fig. S1 - Myotis lucifugus embryonic fibroblast responsiveness to IFNa stimulation.** RT-qPCR showing normalized gene expression for *IFIH1* and *STAT1* in untreated and IFNa-stimulated primary cells. *= paired, two-tailed Student's *t*-test *p*-val < 0.05.

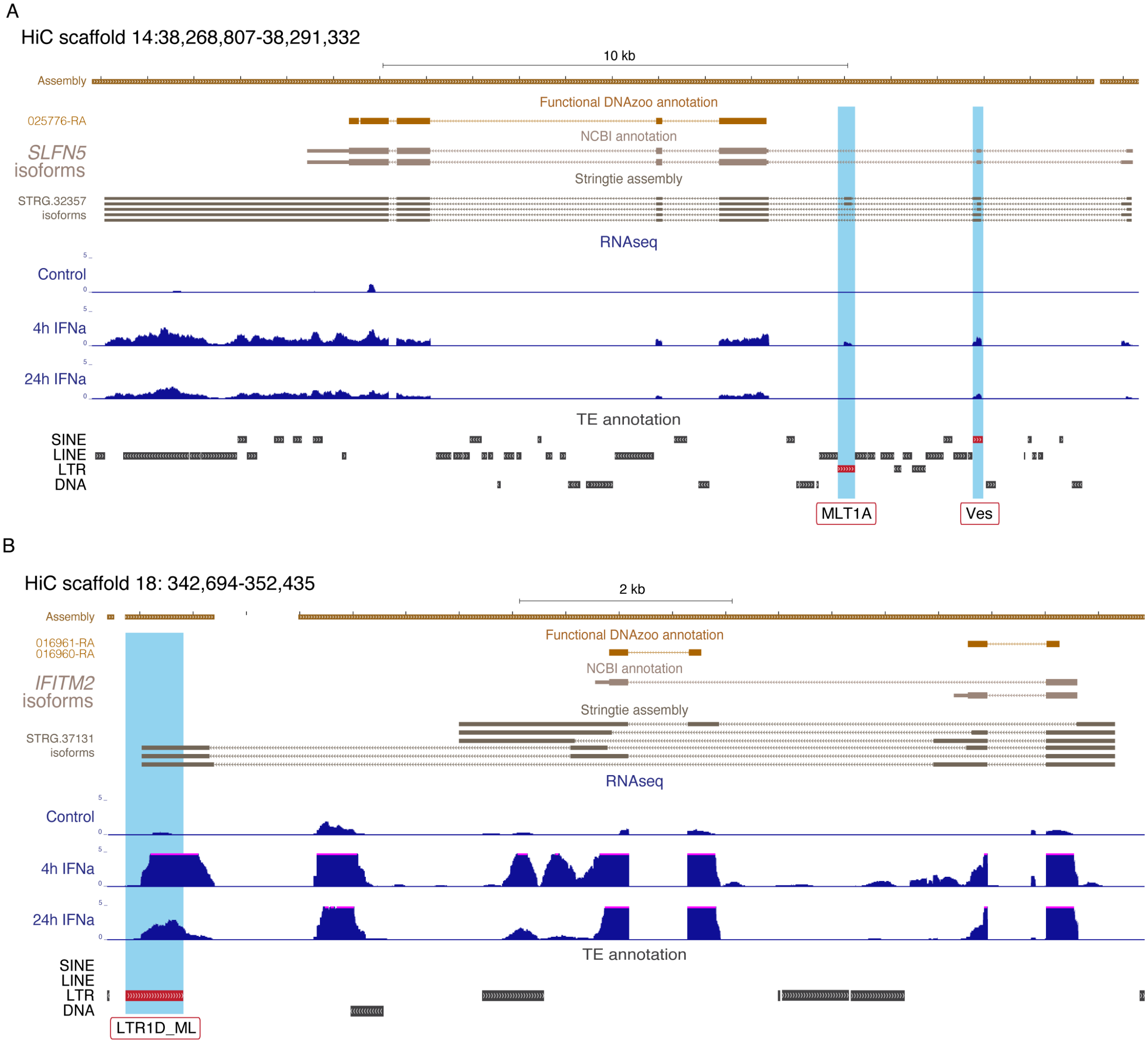


**Fig. S2 - TE exonization events in *Myotis* ISGs. A**) Custom UCSC genome browser screenshot of the SLFN5 locus, where two exons (light blue highlights) represent a potential alternative exon (E3) identified by genome-guided transcriptome assembly in StringTie (MLT1A LTR, on the left) and a constitutive exon (E2; Ves SINE, on the right). RNAseq coverage suggests upregulation of the transcript at 4h post IFNa treatment, consistent with its role in immune responses, and lower expression in 24h post treatment samples compared to unstimulated cells. The alternative isoform that derives from the exonization of the MLT1A LTR may represent an early-response to IFN stimulation specific transcript, as there is no RNAseq coverage signal at 24h post IFN treatment (as for the control) over the alternative E3. **B**) Custom UCSC genome browser screenshot of the IFITM2 locus. Genome-guided transcriptome assembly in StringTie where suggests the presence of a IFN inducible alternative isoform of IFITM2 where the C-terminal region (light blue highlight) derived from the exonization of a Myotis-specific TE (LTR1D_ML). RNAseq coverage suggests upregulation of the transcript at 4h post IFNa treatment, consistent with its role in immune responses, and lower expression in 24h post treatment samples compared to unstimulated cells. Targeted sequencing of the loci or long-read RNA sequencing will be required to validate both IFITM2 and SLFN5 TE-derived isoforms.

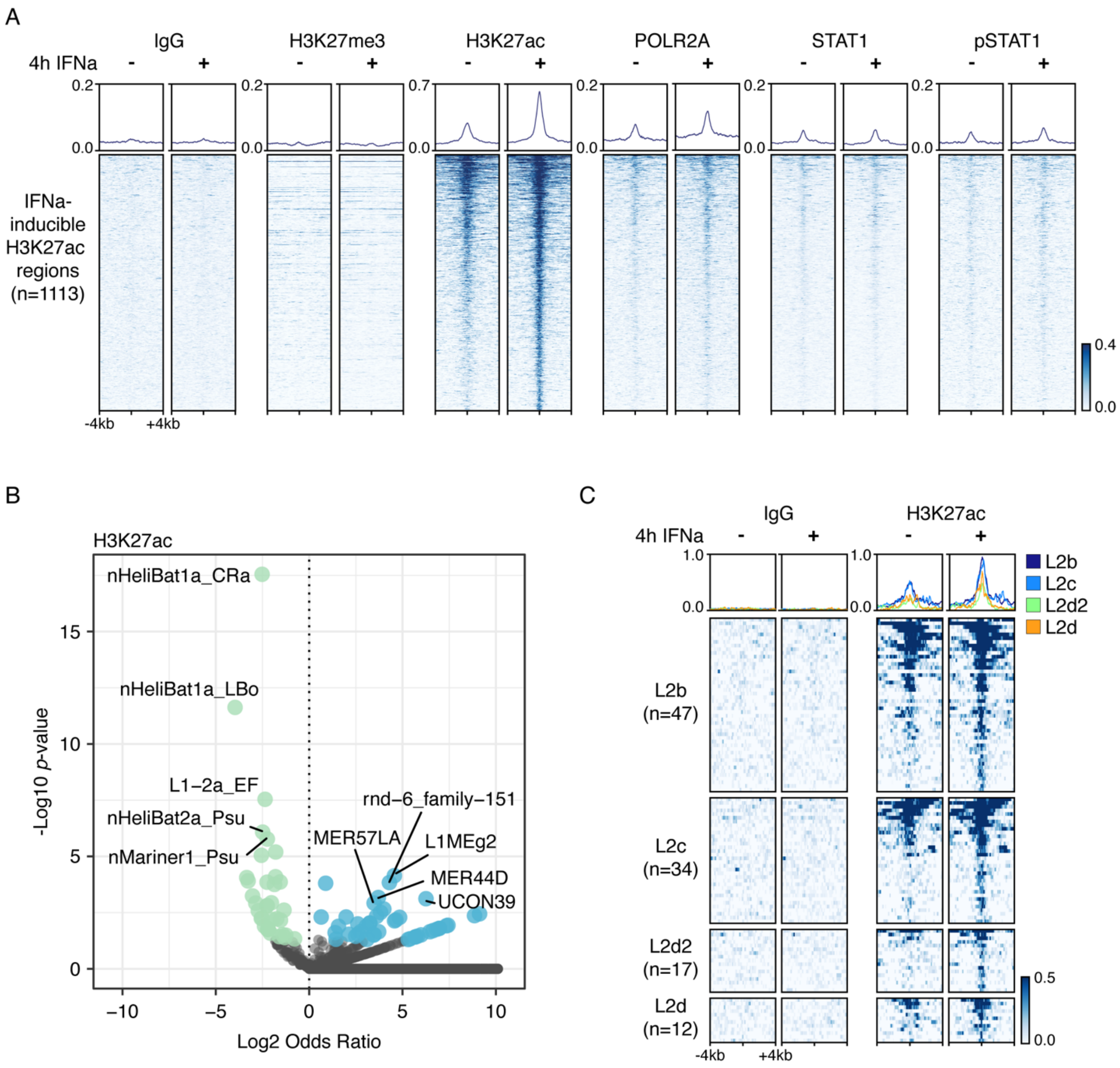


**Fig. S3 -** A) Heatmaps showing normalized CUT&RUN signal (signal per million reads) over IFNa-inducible H3K27ac (unadjusted *p*-val < 0.10, log2FC > 0, n=1113). B) Volcano plot visualizing family-level TE enrichment for IFNa-inducible (unadjusted *p*-val < 0.10, log2FC > 0) H3K27ac regions. TE families with a Fisher's two-tailed *p*-val < 0.05 were defined as enriched (blue) or depleted (green) according to the reported odds ratio. Nonsignificant families are shown in grey. C) Heatmaps showing normalized CUT&RUN signal (signal per million reads) over IFNa-inducible (unadjusted *p*-val < 0.10, log2FC > 0) H3K27ac-marked L2 families.

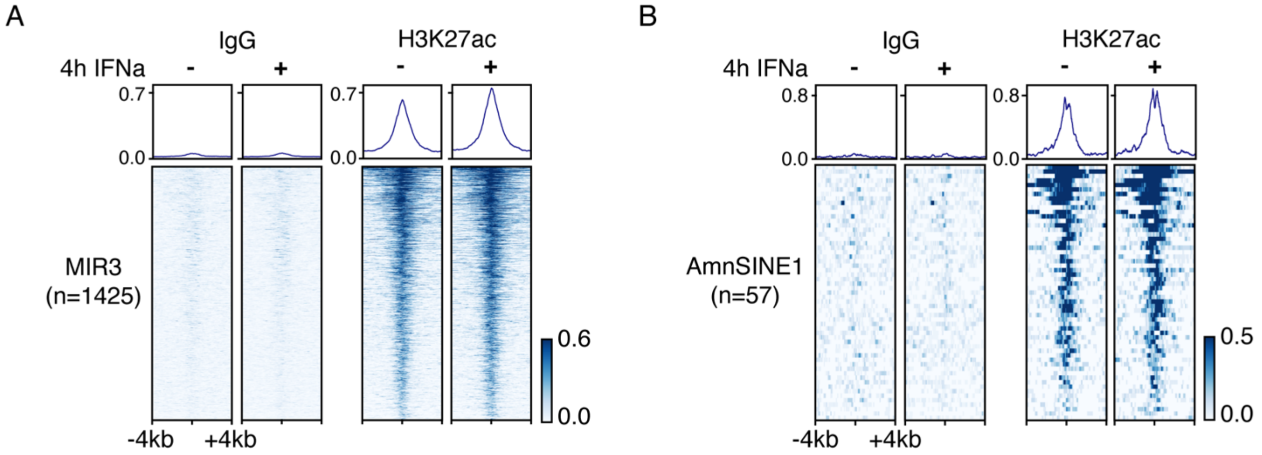


**Fig. S4 -** Heatmaps showing normalized CUT&RUN signal (spike-in signal per million reads) over H3K27ac-marked A) MIR3 and B) AmnSINE1 families.

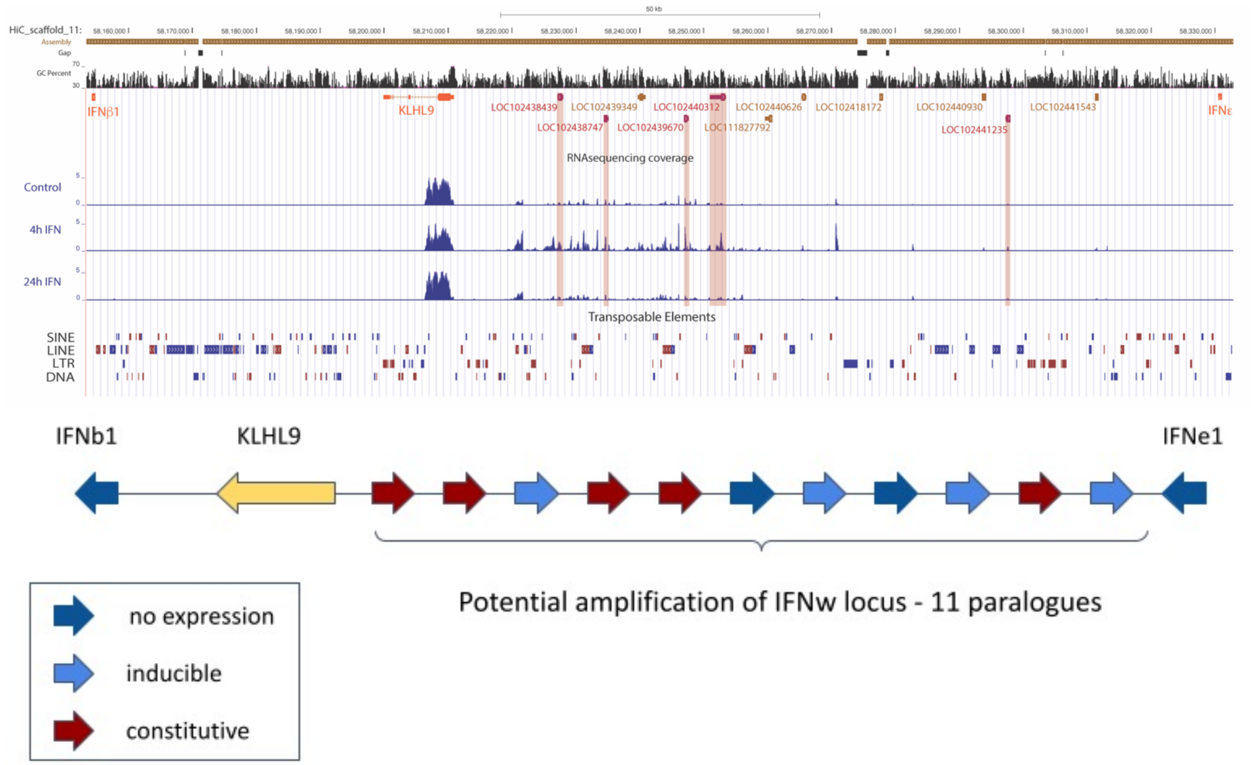


**Fig. S5 - Reconstruction of the type I IFN locus.** On top, custom UCSC genome browser screenshot of the type I IFN locus. We used the IFNb and IFNe1, as well as KLHK9 conserved syntenyc loci as anchors to annotate type I IFN genes. At the bottom, based on previous work (Kepler, T.B., Sample, C., Hudak, K. *et al.* Chiropteran types I and II interferon genes inferred from genome sequencing traces by a statistical gene-family assembler. *BMC Genomics* 11, 444 (2010). doi:10.1186/1471-2164-11-444; Hecker N, Hiller M. A genome alignment of 120 mammals highlights ultraconserved element variability and placenta-associated enhancers. *Gigascience*. 2020;9(1):giz159. doi:10.1093/gigascience/giz159) we were able to identify 11 IFNw paralogue genes. 2 paralogues had no support of being expressed in Myotis embryonic fibroblasts based on RNA-seq expression data (dark blue), 4 paralogues showed no expression in unstimulated cells but induction upon IFN treatment (light blue) and finally 5 paralogues showed basal low expression in unstimulated cells, often followed by mild induction upon IFN treatment.
